# *In silico* study on the effects of disulfide bonds in ORF8 of SARS-CoV-2

**DOI:** 10.1101/2021.11.29.470346

**Authors:** Yadi Cheng, Xubiao Peng

## Abstract

The COVID-19 epidemic, caused by virus SARS-CoV-2, has been a pandemic and threatening everyone’s health in the past two years. In SARS-CoV-2, the accessory protein ORF8 plays an important role in immune modulation. Here we present an *in silico* study on the effects of the disulfide bonds in ORF8, including the effects on the structures, the binding sites and free energy when ORF8 binds to the human leukocyte antigen (HLA-A). Using the explicit solvent Molecular Dynamics (MD) simulations, we collect the conformational ensembles on ORF8 with different disulfide bonds reduction schemes. With a new visualization technique on the local geometry, we analyze the effects of the disulfide bonds on the structure of ORF8. We find that the disulfide bonds have large influences on the loop regions of the surface. Moreover, by performing docking between HLA-A and the conformational ensembles of ORF8, we predict the preferred binding sites and find that most of them are little affected by the disulfide bonds.Further, we estimate the binding free energy between HLA-A and ORF8 with different disulfide bonds reductions. In the end, from the comparison with the available experimental results on the epitopes of ORF8, we validated our binding sites prediction. All the above observations may provide inspirations on inhibitor/drug design against ORF8 based on the binding pathway with HLA-A.

## Introduction

The COVID-19, which is caused by severe acute respiratory syndrome coronavirus 2 (SARS-CoV-2), ^1–4^ has been a public health event all over the world.The virus SARS-CoV-2 encodes 16 non-structural proteins, 4 structural proteins and 6 accessory proteins in total, wherein the open reading frame 8 (ORF8) is an important accessory protein. Different from other proteins, ORF8 in SARS-CoV-2 has very low homology with SARS-CoV,^5–8^ which may explain the higher transmission efficiency of SARS-CoV-2.^9–11^ According to the references,^12–16^ ORF8 in SARS-CoV-2 is a rapidly evolving protein with many mutations and deletions of nucleotides, which may result in two opposite consequences. On one hand, some deletions such as Δ382 may lead to the milder infections of the virus.^17^ On the other hand, the mutations in ORF8 may change the clinical symptoms of COVID-19, which makes the therapies even more difficult. ^18^ In addition, it was found that the variants with a combination of spike mutations and the truncation of ORF8 may be more spreading. ^19^

As an accessory protein, ORF8 is compose of 121 amino acids with a signal sequence (residues 1-15) located in the N-terminal for invading the endoplasmic reticulum (ER),^9,20,21^ where the protein ORF8 can interact with a series of host proteins. Studies have shown that ORF8 in SAR-CoV-2 can down-regulate the expression of the major histocompatibility complex class I (MHC-I) proteins to mediate the immune escape.^11^ For human, the human leukocyte antigen (HLA-A) is an important protein belonging to MHC-I. Hence, it is important to study the mechanism on the binding between ORF8 and HLA-A for developing new therapy against SRS-CoV-2. Up to now, there have been several experimental and theoretical studies on identifying the epitopes on ORF8.^22–25^

Structurally, the monomeric ORF8 in SARS-CoV-2 contains three disulfide bonds (ss-bond) as shown in Fig 1. In reference 20 it was pointed out that ORF8 can have two different dimeric states, known as the covalent dimer and non-covalent dimer. The covalent dimer is formed by an intermolecular disulfide bond between residues C20 in the N-terminals of both monomers as labeled in Fig 1, while the non-covalent dimer is stabilized by the segment ^73^YIDI^76^ on the interface of the non-covalent dimer. In this paper, we only focus on the former for the dimer case. We notice that the residues for forming the disulfide bonds seems to be conserved although there have been many mutations in ORF8.*In vivo*, the correct formation of the disulfide bonds is catalyzed by the protein disulfide isomerase (PDI). However, it was found that the red wine and grape juice can inhibit the activity of PDI, which may lead to the loss of the disulfide bonds of the proteins,^26^ such as ORF8.

**Figure 1:**
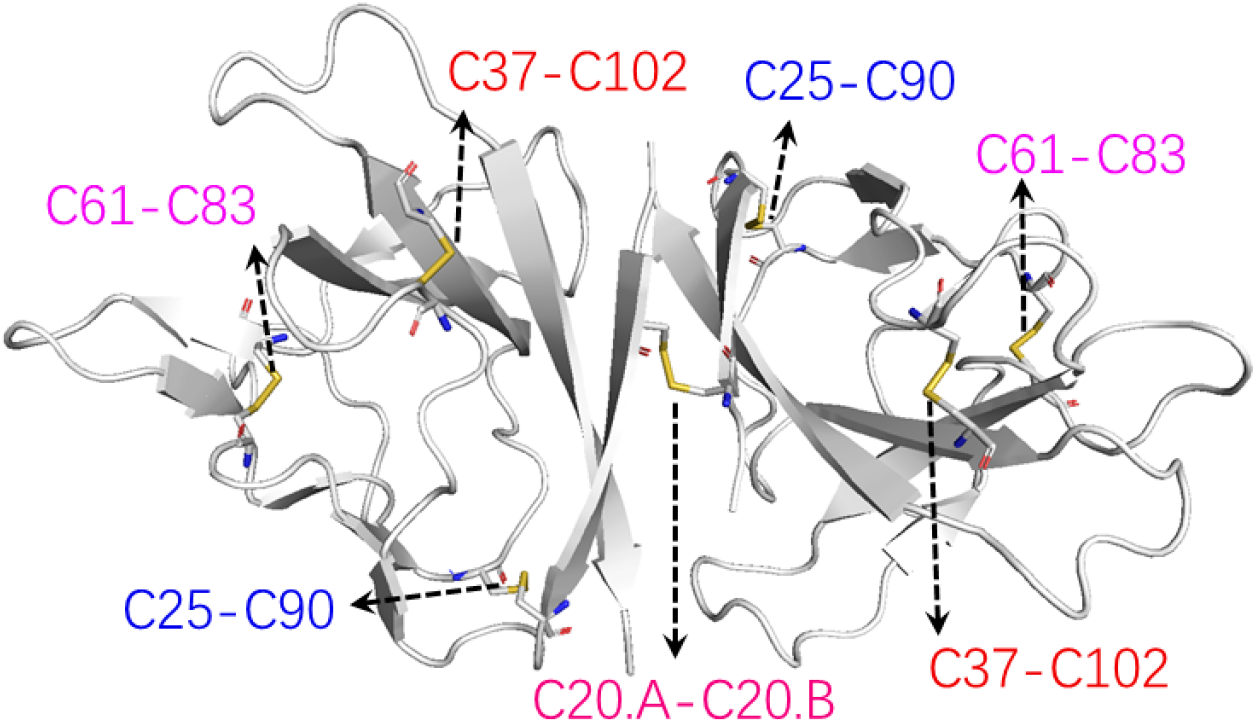
The disulfide bonds in the structure of a covalent-bonded dimeric ORF8. The residues forming disulfide bonds are labeled. The structure is taken from PDB entry 7JX6.

In this article, we investigate the influence of the disulfide bonds on the structure and the binding properties of ORF8 using *in silico* methods including the molecular dynamics (MD) simulations and docking. From the MD simulations on ORF8 with different disulfide bonds reduced, we find the most affected residues/segments through a new visualization technique based on the Discrete Frenet Frame (DFF).^27–29^ Based on the conformational ensembles sampled from MD simulations, we further investigate the influences of the disulfide bonds on the binding sites between ORF8 and HLA-A using the docking method. In the end, we compared the binding affinities between ORF8 and HLA-A when different disulfide bonds in ORF8 get reduced. We find that the broken of the disulfide bonds leads to large changes on local conformations, but does not have essential influences on the binding properties with HLA-A.

## Materials and Methods

### MD simulation

We use the software package GROMACS 5.1.1^30^ for the MD simulations, with force field CHARMM36M^31^ and the modified TIP3P^32^ water model which is adjusted for CHARMM force field. In the simulation, the LINCS algorithm is used for the covalent bonds connected with hydrogen atoms, and the time step is set to 2 fs. The protein is solvated in 0.15 mmol/L KCl solution in order to model the intracellular environment. After a short energy minimization on the system, we perform an NVT simulation for 100 ps with the V-rescale temperature coupling at 300 K, followed by an NPT simulation for 300ps with the Parrinello-Rahman coupling method at reference pressure 1bar. The relaxation time for temperature coupling and pressure coupling are 0.1 ps and 2ps, respectively. In the end, the production simulations are performed for 200ns at 300 K for generating the conformational ensembles.

### Local geometry visualization

In order to find the residues that are most affected by the disulfide bonds reductions in ORF8, we employ the visualization method described in the references,^27–29^ where we introduced the Discrete Frenet Framework (DFF) to describe the *C_α_* trace of a protein chain. As shown in Fig 2(A), the virtual bond angle *κ*_*i*+1_ is determined by three consecutive *C_α_* atoms (**r**_*i*_,**r**_*i*+1_,**r**_*i*+2_), and the virtual torsional angle *τ*_*i*+1_ is determined by four *C_α_* atoms (**r**_*i*–1_,**r**_*i*_,**r**_*i*+1_,**r**_*i*+2_). The angle pairs (*κ,τ*) can be visualized by stereographical projection as described in reference^29^ and Fig 2(B), where the standard (*κ,τ*) distributions for different secondary structures are also indicated.

**Figure 2:**
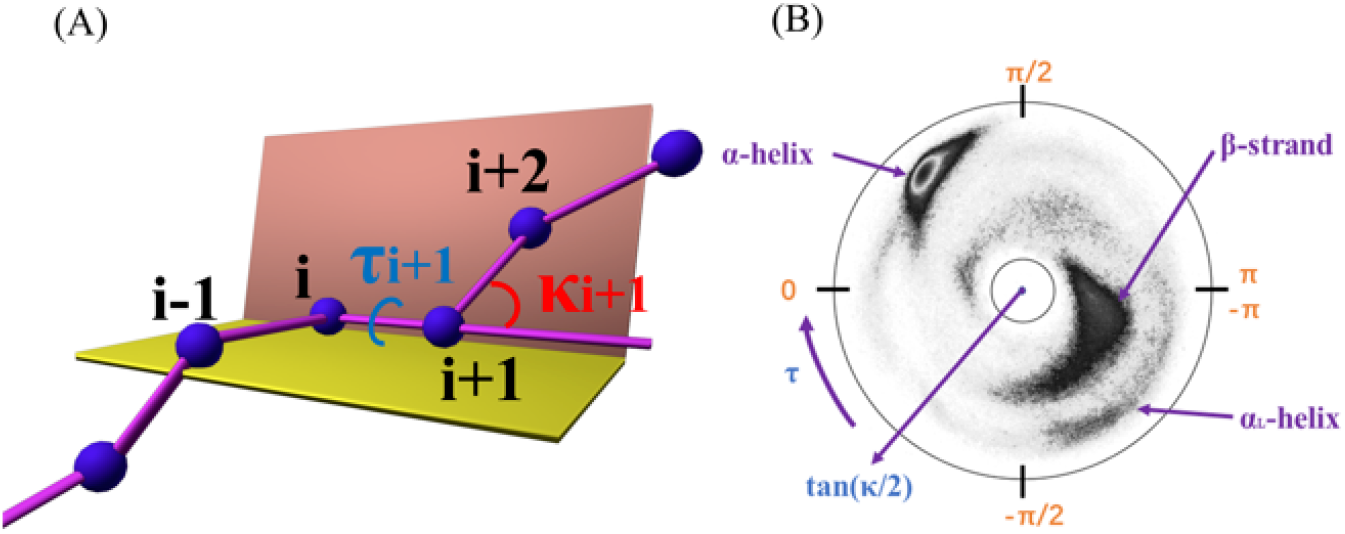
(A) Definitions of virtual bond (*κ*) and torsion angles(*τ*).(B)The stereographical projection map of (*κ,τ*). The secondary structures such as *α*-helix, *β*-strand and *α_L_*-helix have been labelled in the distribution.

### Binding sites prediction

On the binding site prediction, we use the webserver ZDOCK^33^(https://zdock.umassmed.edu/) to predict the binding sites between ORF8 and HLA-A. The webserver ZDOCK is a docking program based on fast Fourier transform and 3D convolution library, where the proteins are assumed to be rigid. It searches all the possible mutual binding modes obtained by relative translations and rotations between two proteins, and then ranks them according to an energy-based scoring function, which takes the potential energy, spatial complementarity and electric field force into account. The mode with a higher score indicates that it has a lower potential energy, and hence is also ranked higher. In this paper, we selected the top mode with the highest score. With the obtained binding mode, we further analyze the hydrogen bonds and salt bridges between ORF8 and HLA-A with the PDBe-PISA webserver^34–36^ (http://pdbe.org/pisa/), which can directly give the possible hydrogen bonds and salt bridges between the two proteins in the complex. Hence, for a particular configuration of ORF8, we can obtain its binding sites and interactions with HLA-A determinately through the above workflow.

For each state of ORF8 with different disulfide bonding in our study, we sample 800 configurations along the trajectory from the MD simulations, repeat the above workflow for each configuration, and finally get the distributions of the possible binding sites and interactions.

### Binding free energy estimation

Starting from the representative configurations of the complex predicted from ZDOCK, we perform MD simulations for 200ns to observe the stabilities of the complexes. The final curve of the binding free energy between ORF8 and HLA-A changing with simulation time are evaluated along the trajectory using the webserver PRODIGY^37,38^ (https://bianca.science.uu.nl/prodigy/), where the binding free energy is roughly estimated based on the number of interfacial contacts, according to the observation that the number of interfacial contacts in the complex correlates with the experimental binding affinity^39^ well.

## Results and Discussion

The protein ORF8 can be in either monomeric or dimeric state. However, the only structures for ORF8 deposited in PDB (PDB code: 7JX6 or 7JTL)^20^ are in the dimeric state with the intermolecular disulfide bond formed, which we denoted as ss-dim state throughout the paper. To investigate the influences of the disulfide bonds, we further prepare four additional states by reducing different types of the disulfide bonds from the native ss-dim state, including the dimeric state with *only the intermolecular* disulfide bond reduced, the dimeric state with *all* the disulfide bonds reduced, and the monomeric states *with and without all the intramolecular* disulfide bonds, which are denoted as int-sh-dim, all-sh-dim, ss-mon and sh-mon states, respectively.

For the MD simulation, we choose the PDB entry 7JX6 as the initial conformation since it has higher resolution. However, we notice that the three-dimensional structure for the segment 63-78 in chain B is missing. As a result, we reconstruct the missing structure using the loop refinement software MODELLER^40–43^ before the simulations on dimeric states. For the simulations on monomer, we simply take chain A as the initial configuration. The peptide chain of ORF8 starts from residue Q18, which is due to the fact that the signal sequence in N-terminal has been cleaved when invading ER. Hence, we cap the N and C termini of the PDB entry with NH3+ and COO-, respectively.

For each of the five states, we perform the MD simulations for 200ns, from which only the last 160ns is used for collecting conformations. Each simulation is replicated three times to ensure the reproducibility and convergence. The all-atom root mean square deviation (RMSD) for the three replicated MD simulations on five states are shown in Fig S1 in the Supplemental information, indicating that the simulations get converged in the end.

### The influence of the disulfide bonds on the global structures

The radius of gyration (Rg) is usually used to characterize the compactness of the protein. A smaller Rg usually indicates a more compact structure. By comparing the evolutions of the Rg values shown in Fig S2, we can see that the disulfide bonds can increase the compactness of the protein for both monomer and dimer. In Fig 3, we show the distributions of the solvent accessible surface area (SASA) for the simulations of five states. For the dimeric states, the SASA in the ss-dim state is slightly larger than in the “int-sh-dim” state, and both are smaller than that in the all-sh-dim state. Hence, the effect of the intramolecular disulfide bonds is much more obvious than the intermolecular disulfide bond for dimer. For monomer, the SASA in the sh-mon state is just slightly larger than in ss-mon state due to the loss of the intramolecular disulfide bonds. In the end, we notice that the SASA values in all the dimeric states are much smaller than the twice of the monomeric states, indicating that the dimer is not split into two monomers even with all the disulfide bonds broken. In Fig S3, we show the distributions of the buried surface area for the three dimeric states.

**Figure 3:**
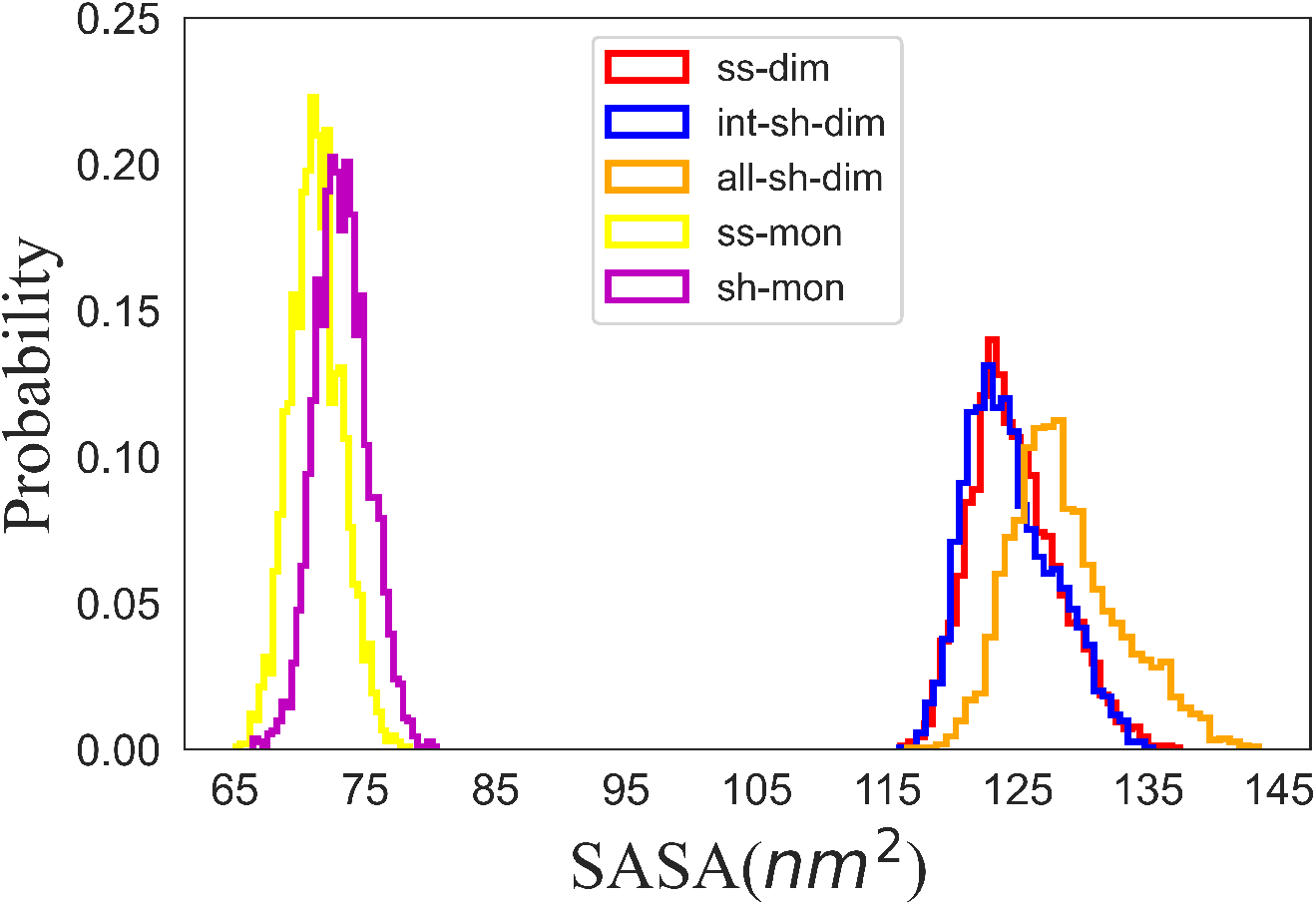
The distributions of the solvent accessible surface area (SASA)for different states.

### The influence of the disulfide bonds on the local structures

With the help of the visualization technique described in the section Local geometry visualization in Material and methods and references,^27–29^ we compared the local geometrical properties of ORF8 for the five states. The local geometrical properties are characterized by the distributions of (*κ_i_,τ_i_*) angles obtained from the conformational ensembles, which are collected from all the three replicated MD simulations for each state.

As an example, we show the comparison of(*κ,τ*) distributions for the sequence ^73^YIDI^76^, which is an important segment for stabilizing the non-covalent dimer as shown in reference^20^. In Fig 4, we can see that the distributions are quite different among the simulations of the five states. For all the five states, residues Y73 and I76 are most affected by disulfide bonds reductions, while I74 and D75 are the least affected. For dimers, the widest and most localized distributions of (*κ,τ*) are corresponding to the all-sh-dim and int-sh-dim state, respectively. We can see that the reduction of the intermolecular disulfide bond (from the comparison between “ss-dim” and “int-sh-dim”) decreases the flexibility while further reduction of the intramolecular disulfide bonds (see the comparison between “int-sh-dim” and “all-sh-dim”) increases the flexibility of segment 73-76. However, for monomer it seems that the distributions in sh-mon state are more localized than in ss-mon state, indicating that the reduction of the disulfide bonds decreases the flexibility of this segment when in monomeric state. Hence, we can see that the same disulfide bonds reduction affects the local segment differently, depending on whether the system is in monomeric or dimeric states.

**Figure 4:**
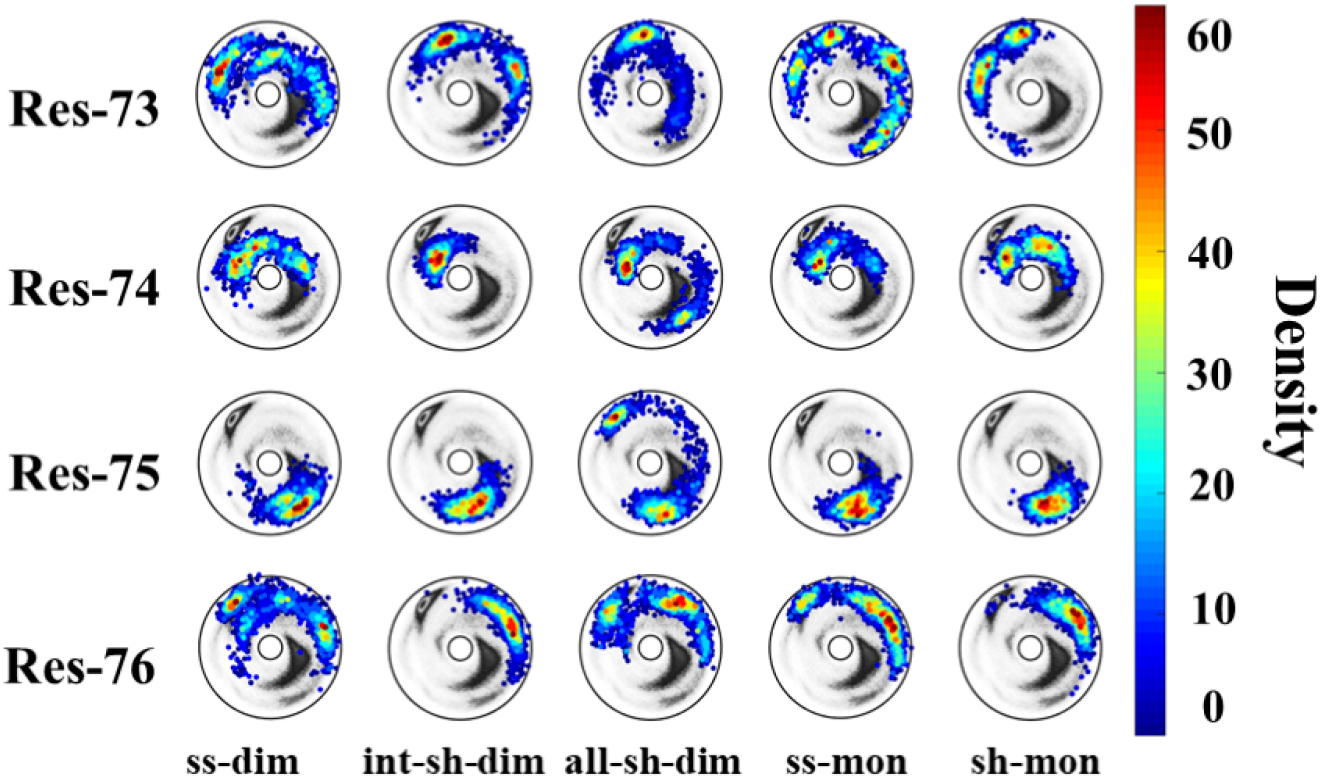
The distributions of the angle pairs (*κ_i_*,*τ_i_*) with *i* ∈ [73, 76] for the segment ^73^YIDI^76^.

By going along the protein chain and comparing the distributions of corresponding (*κ,τ*) angle at each site for the five states, we list all the residues that most affected by different disulfide bonds reductions in Table 1. In Fig S4 we mark all the highly affected residues and find that most of them are located in the loop regions on the surface of the protein.

**Table 1:**
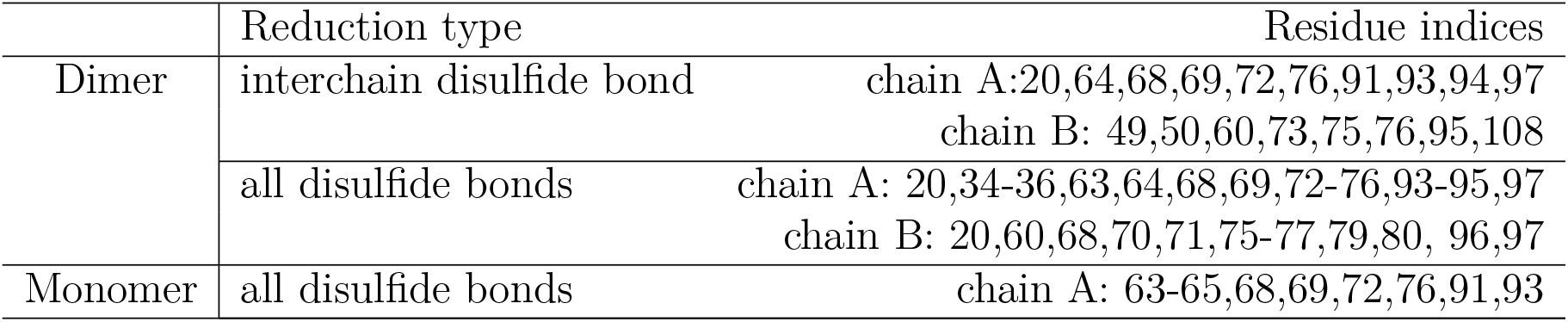
The residues with the conformations most affected by the disulfide bonds reductions.

|  | Reduction type | Residue indices |
| --- | --- | --- |
| Dimer | interchain disulfide bond | chain A: 20, 64, 68, 69, 72, 76, 91, 93, 94, 97<br>chain B: 49, 50, 60, 73, 75, 76, 95, 108 |
|  | all disulfide bonds | chain A: 20, 34-36, 63, 64, 68, 69, 72-76, 93-95, 97<br>chain B: 20, 60, 68, 70, 71, 75-77, 79, 80, 96, 97 |
| Monomer | all disulfide bonds | chain A: 63-65, 68, 69, 72, 76, 91, 93 |

In addition, we find some segments whose local structures are rather conserved across the different simulations. In Fig S5, Fig S6 and Fig S7, we show the distributions of (*κ,τ*) angles for segment 39-42, 104-107 and 110-112 in three replicated MD simulations, respectively. We can see that the three segments have very conserved patterns of the distribution, which is independent on the status of the disulfide bonds and the chains.

### Influence of the disulfide bonds on the binding properties between ORF8 and HLA-A

From each of the simulations on ORF8 with different states, we sample 800 conformations along the trajectory with the first 40ns discarded. For HLA-A, we take the entry with PDB code 3HLA^44^ as the input structure. Following the method described in subsection Binding sites prediction in Section “Material and methods”, we analyze the interactions between ORF8 and HLA-A when binding happens.

The interactions are classified into two classes named as hydrogen bonds and salt bridges. In Fig 5(A) and (B) we show the distributions of the binding sites of ORF8 with hydrogen bonds on the interface of the complex in different states. We note that here we only show the binding sites with probability larger than 10% in Fig 5. We can see that most of the preferred sites in Fig 5 are not much affected by the disulfide bonds reduction, such as the sites 40, 42, 104, 105, 106, 110 in chain A and sites 39, 40, 105, 110 in chain B. However, different states can also have their specific binding sites for forming hydrogen bonds. For example, in ss-dim the probability of site 19 in chain A is relatively higher; In int-sh-dim, site 18 in chain A is outstandingly high; And in ss-mon, sites 41 and 43 in chain A are outstandingly high. In addition, we also find that the chain A of ORF8 is more likely to bind with HLA-A in hydrogen bonds than the chain B. In the Supplemental information, we also show the hydrogen bonding sites of protein HLA-A in Fig S8. In HLA-A, the sites 65 and 155 of chain A are the most preferred hydrogen binding sites.

**Figure 5:**
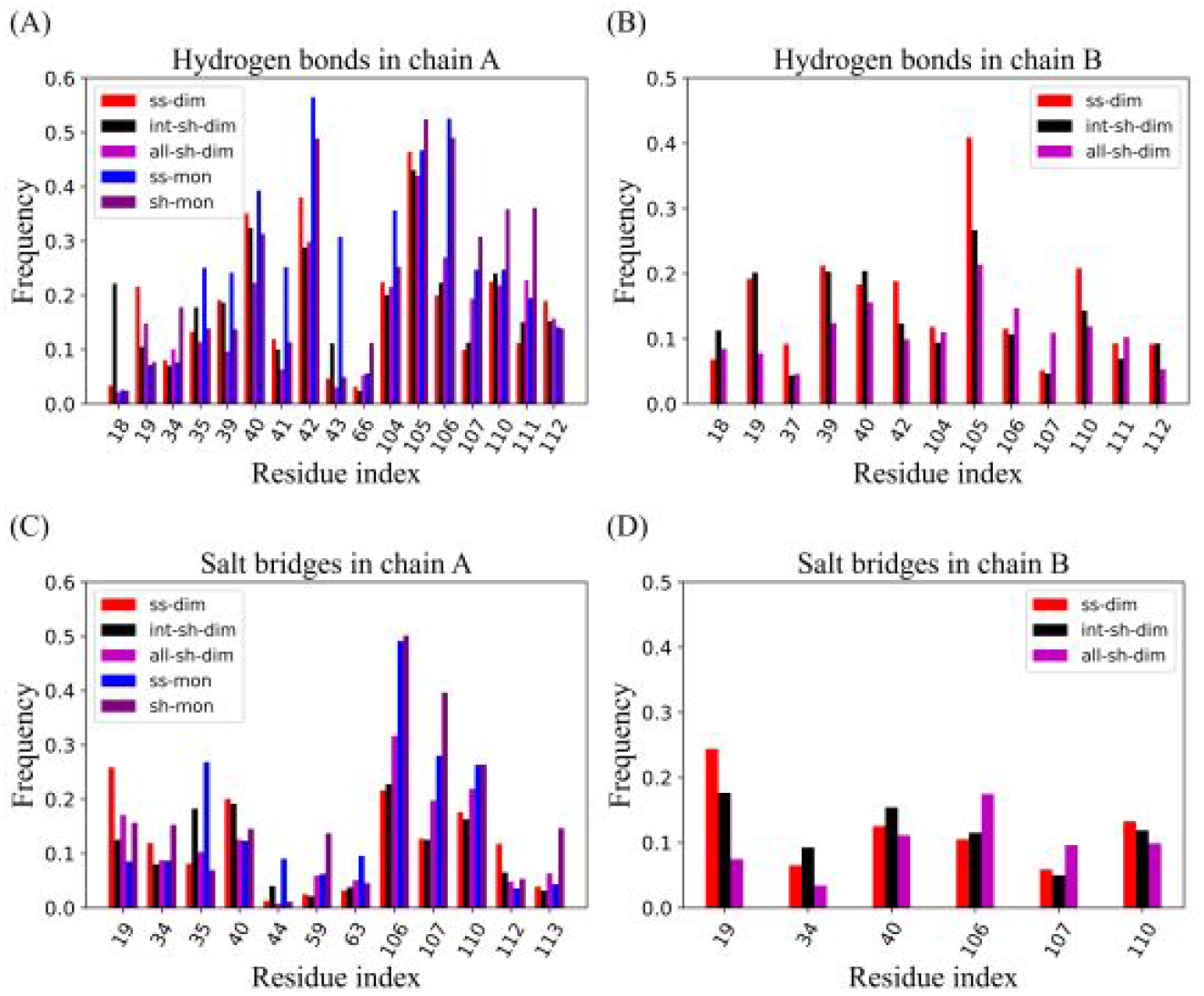
The distributions of the binding sites formed in hydrogen bond with HLA-A for (A) chain A and (B) chain B of ORF8.The distributions of the binding sites formed in salt bridges with HLA-A for (C) chain A and (D) chain B of ORF8.

Similarly, we show the distributions of the residues on ORF8 binding with HLA-A in salt bridges in Fig 5(C) and (D). We find that the preferred binding sites in common across the five states are 19, 40, 106,107,110 in chain A. Some salt bridges that are specific for certain ORF8 states are listed as follows: For ss-dim state, site 19 is relatively higher; For int-sh-dim and ss-mon states, site 35 is higher; For sh-mon state, site 59 and site 113 are higher. We also draw the distributions of the salt bridge binding sites on HLA-A in Fig S9, where the sites 60, 66,146 in chain A are the hot spots on HLA-A for binding with ORF8 across the different states.

In summary, we can see that most of the binding sites with high probabilities are independent on the status of the disulfide bonds reductions and they are mainly located in chains A for both ORF8 and HLA-A. In particular, we notice that the common binding sites 40, 42, 104,105, 106, 107 are well consistent with in the conserved segments 39-42 and 104-107 that are found in the previous subsection The influence of the disulfide bonds on the local structures.

### The influence of the disulfide bonds on the binding stability between ORF8 and HLA-A

In order to explore the influences of the disulfide bonds on the stability of complex formed between ORF8 and HLA-A, we selected a representative structure as the initial configuration to perform MD simulation for each of the five disulfide bonding states. We require that the representative complex must follow the main property of the distributions as shown in Fig 5(A), Fig S8, Fig 5(C) and Fig S9. In Fig S10,we show the 3D structures of the representative complex with the corresponding hydrogen bonds and salt bridges labelled. The RMSD curves for the five MD simulations are shown in Fig S11, indicating that the systems are all well equilibrated.

In Fig 6 we show the number of hydrogen bonds on the interface between ORF8 and HLA-A during the MD simulations for each complex. We can see that the number of hydrogen bonds gets converged along each trajectory. For the dimers, it seems that the hydrogen bonds have the largest number in the ss-dim state, followed by the “all-sh-dimer” state, and it has the smallest number in the int-sh-dim state. For monomer, sh-mon has more hydrogen bonds than ss-mon. In the end, the binding free energy for each complex is estimated along the trajectory using PRODIGY webserver^37,38^ and the evolution curves with simulation time are shown in Fig 7. We can see that the ss-dim and all-sh-dim states have the lowest free energy and hence are the most stable. In contrast, the ss-mon state has the largest free energy and is the most unstable one when forming complex with HLA-A. We note that such a result is consistent with the hydrogen bonds analysis.

**Figure 6:**
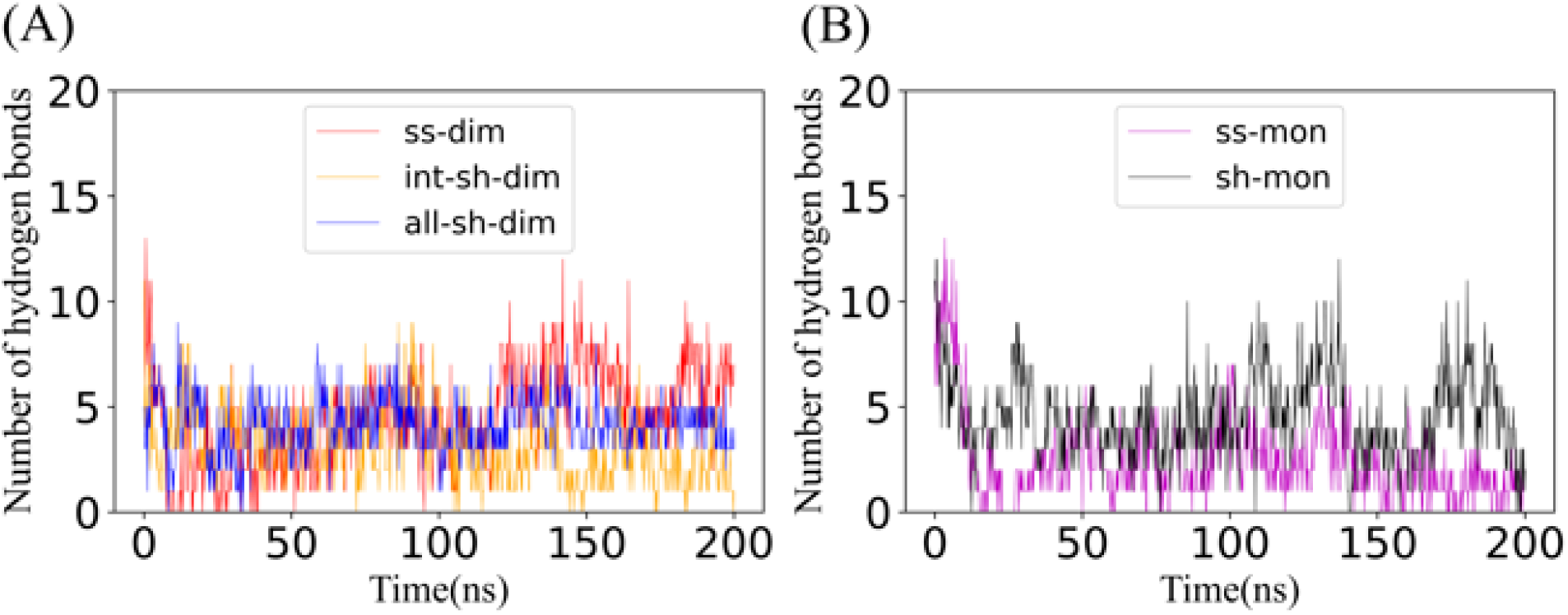
The number of hydrogen bonds between HLA and ORF8 in different states. (A)The three dimeric states. (B) The two monomeric states.

**Figure 7:**
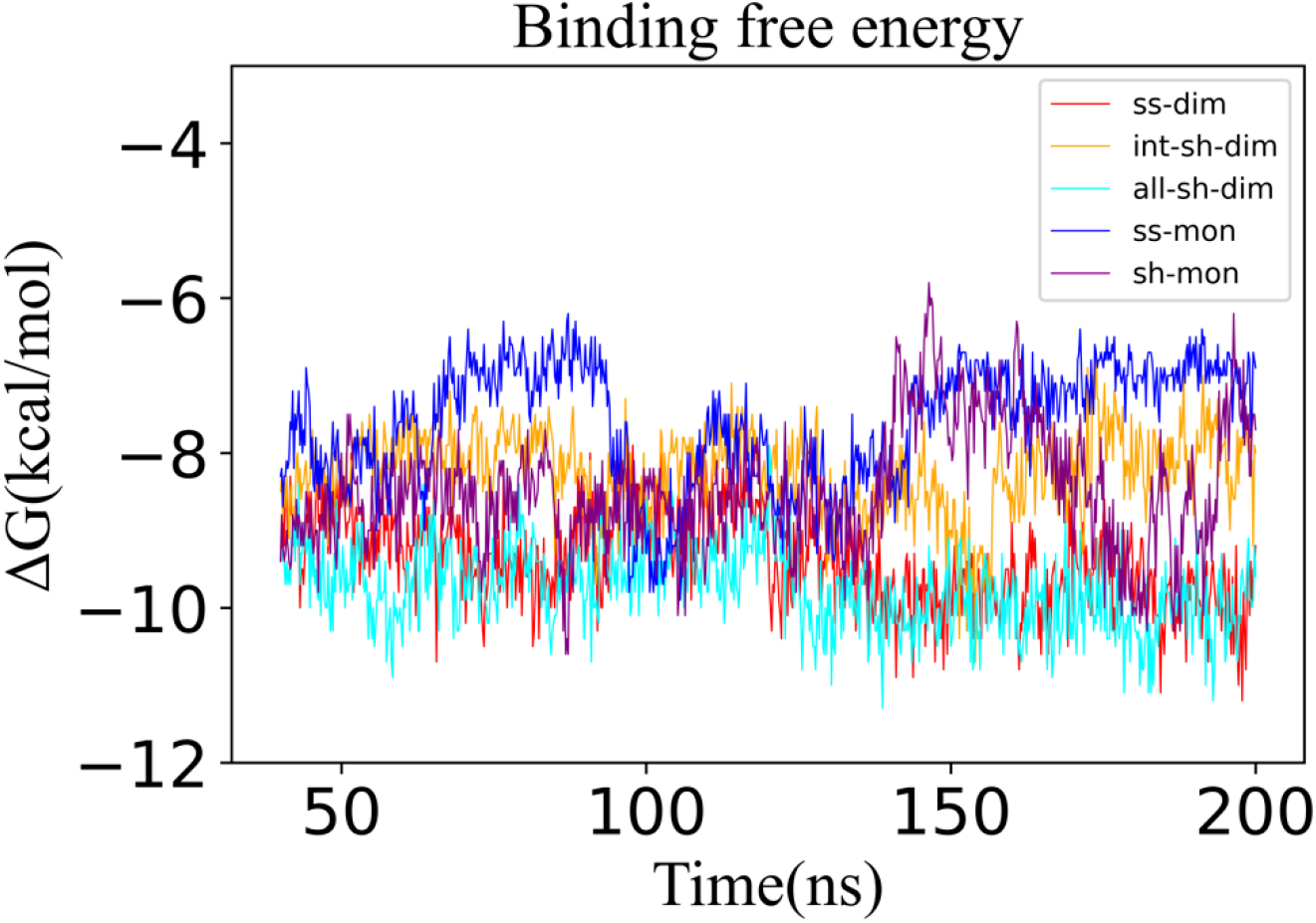
The evolution curves of the binding free energy for the five docking complexes in MD simulations.

### Binding Sites Validation

We compared our prediction on the preferred binding sites in ORF8 with the experimental studies on the epitope identification as shown in Fig 8. We can see that the three consensus binding sites 39-43(IHFYS),104-107(FYED) and 110-112(EYH). The segment 39-43(IHFYS) is well/partially overlapped with the identified epitopes in references 23-25. The segments 104-107(FYED) and 110-112(EYH) in our prediction are corresponding to the epitope106-110(EDFLE) in the literature 22. We note that all these binding sites are little affected by the disulfide bonds according to our local geometrical analysis. (See Fig S5, Fig S6 and Fig S7). Such conserved binding sites can be helpful in designing inhibitors against ORF8 of SARS-CoV-2.

**Figure 8:**
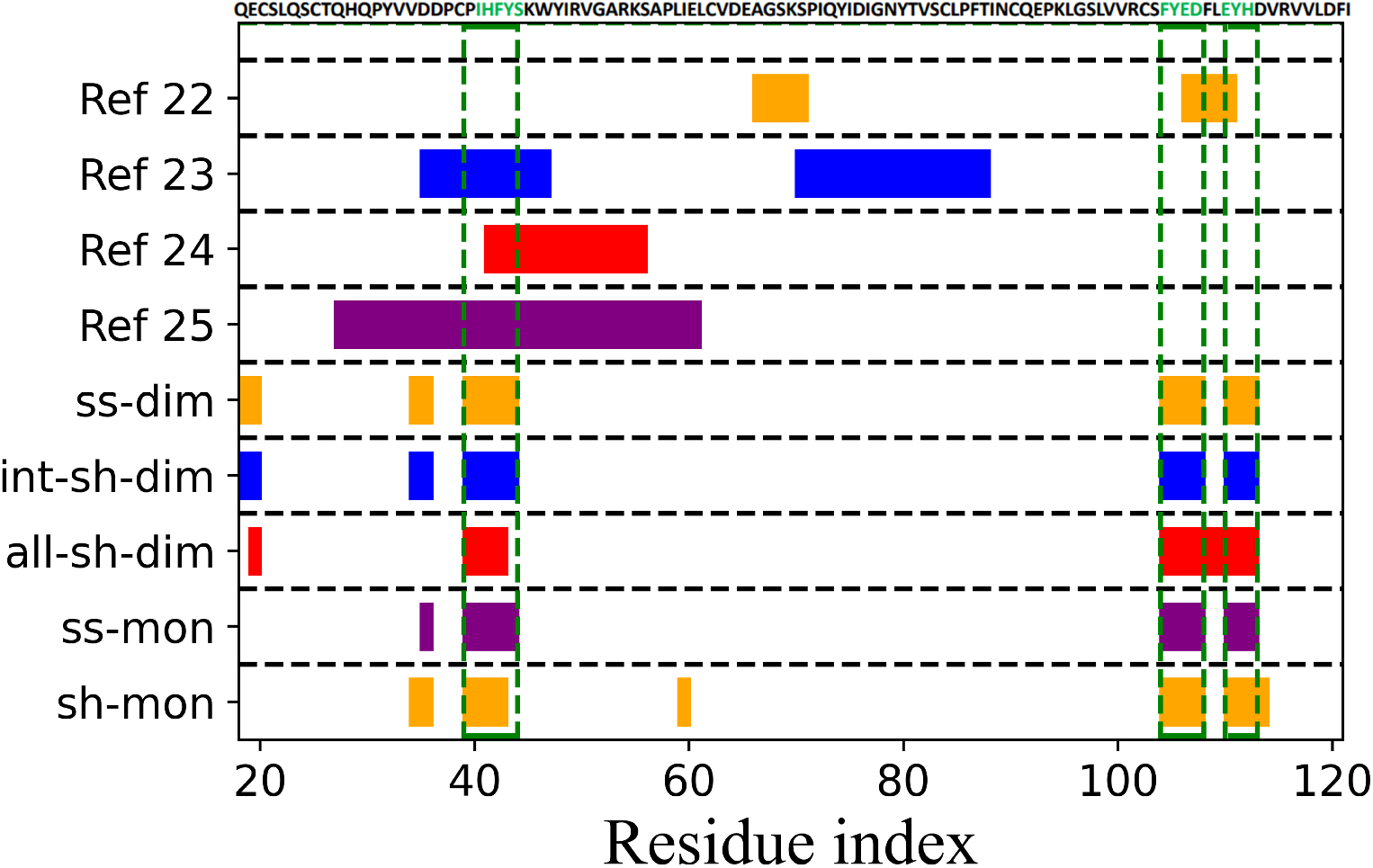
The comparison between our prediction results and the results in literatures. The green boxes are the consensus binding sites with different disulfide bond reduction schemes. We also colored these binding sites in green in the sequence. The results listed in lines 1-3 are the epitopes identified from the experimental literatures 22-24 while the result in line 4 are the predicted epitopes by literature 25. The last five lines are our predicted binding sites with different disulfide bond reduction schemes.

## Conclusion

In this article, we investigated the influences of the disulfide bonds on the structure and binding properties of ORF8 through MD simulations on its five states, which are obtained through different disulfide bonds reduction schemes from the native covalent dimeric structure deposited in PDB. By comparing the structural properties of ORF8 at different states, we find the segments that are the most and least affected by the disulfide bonds, respectively. Furthermore, we compared the binding sites between HLA-A and different states of ORF8. We find that the most preferred binding sites are mainly located in segments 39-42, 104-107 and 110-112, partially consistent with the epitope prediction experiments.^22–25^ Since both segments are little affected by the disulfide bonds, they may be good target for further designing antibodies/drugs. By further MD simulations on the complexes formed by ORF8 and HLA-A, we confirm their stabilities and find that the ORF8 in ss-dim and all-sh-dim states can bind HLA-A better than other states.

Based on the above observations, we conclude that the reductions of the disulfide bonds do not greatly affect the binding sites between ORF8 and HLA-A, which is out of our expectation. However, such a property can be helpful on developing drugs against ORF8. For example, one can try to develop inhibitors to prevent ORF8 from binding with HLA-A, so that the immune system can recognize the virus. In the end, we note that our current research is based on the structures of wild type ORF8. For the effective drug design, the effects of different mutants should be taken into further consideration.

## Author Contributions

Xubiao Peng designed the project. Yadi Cheng carried out all the simulations. Yadi Cheng and Xubiao Peng analyzed the data and wrote the article together.

## Acknowledgement

The authors thanks the financial support provided by the Young Scholars Fund of Beijing Institute of Technology.

