## Supplementary information for "*In silico* study on the effects of disulfide bonds in ORF8 of SARS-CoV-2"

**Contents:**

**Figure S1**: The curves of RMSD evolution for the three replicated MD simulations on five different disulfide bonding states.

**Figure S2**: The Rg curves for the three replicated MD simulations on five different disulfide bonding states.

**Figure S3**: The distributions of the buried surface area for the three dimeric states.

**Figure S4**: The locations of the residues that most affected by different disulfide bonds reduction schemes.

**Figure S5**: The distributions of the angle pairs ($\kappa_i$ ,$\tau_i$) with $i \in [39, 42]$ for the segment $^{39}$IHFY$^{42}$.

**Figure S6**: The distributions of the angle pairs ($\kappa_i$ ,$\tau_i$) with $i \in [104, 107]$ for the segment $^{104}$FYED$^{107}$.

**Figure S7**: The distributions of the angle pairs ($\kappa_i$ ,$\tau_i$) with $i \in [110, 112]$ for the segment $^{110}$EYH$^{112}$.

**Figure S8**: The distributions of the binding sites on HLA-A forming in hydrogen bonds with ORF8.

**Figure S9**: The distributions of the binding sites on HLA-A forming in salt bridges with ORF8.

**Figure S10**: Five representative structures for the complex formed by ORF8 binding with HLA-A.

**Figure S11**： The curves of the RMSD evolution for the MD simulations on the five representative complexes.

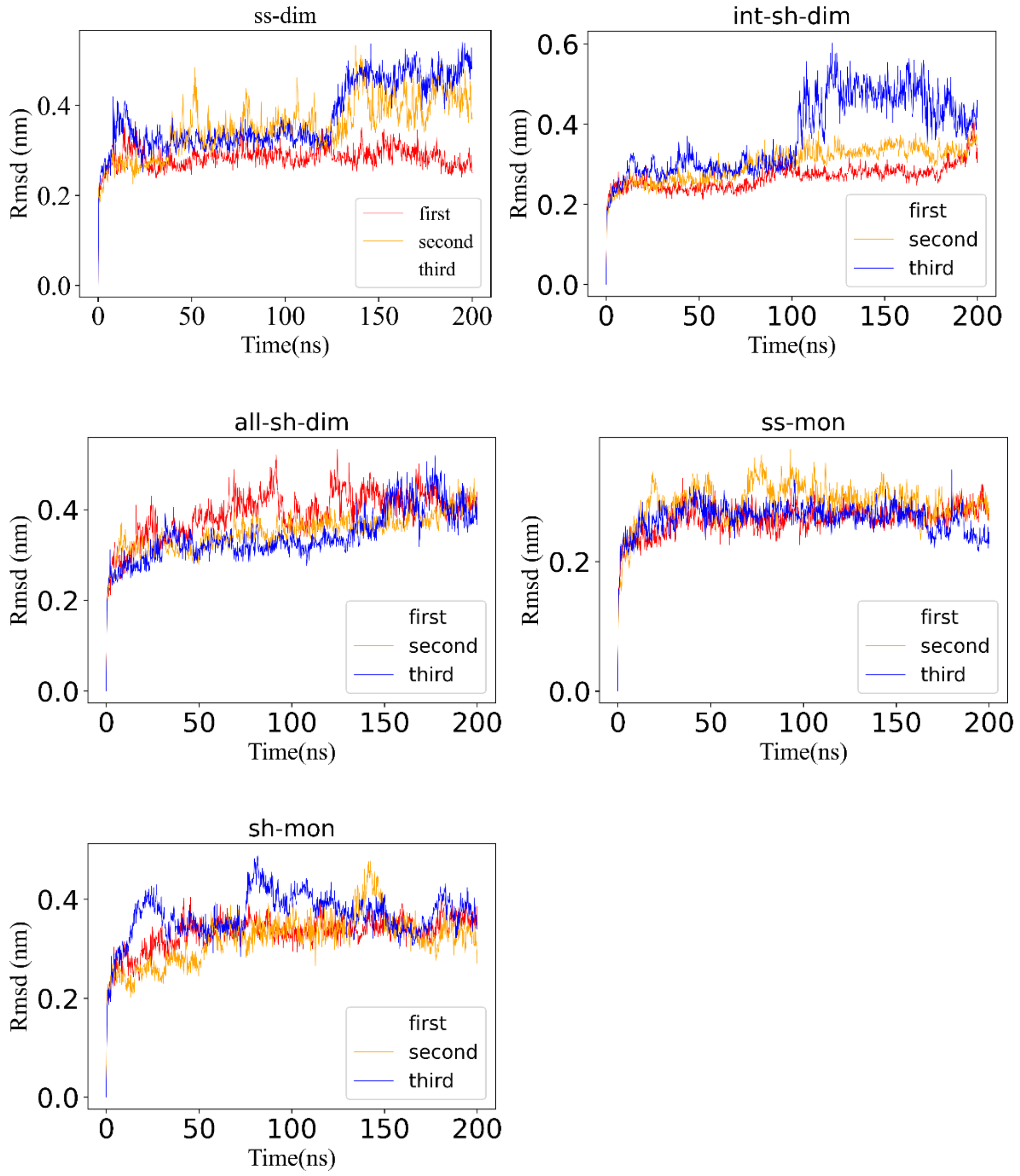


Figure S1 The curves of RMSD evolution for the three replicated MD simulations on five different disulfide bonding states.

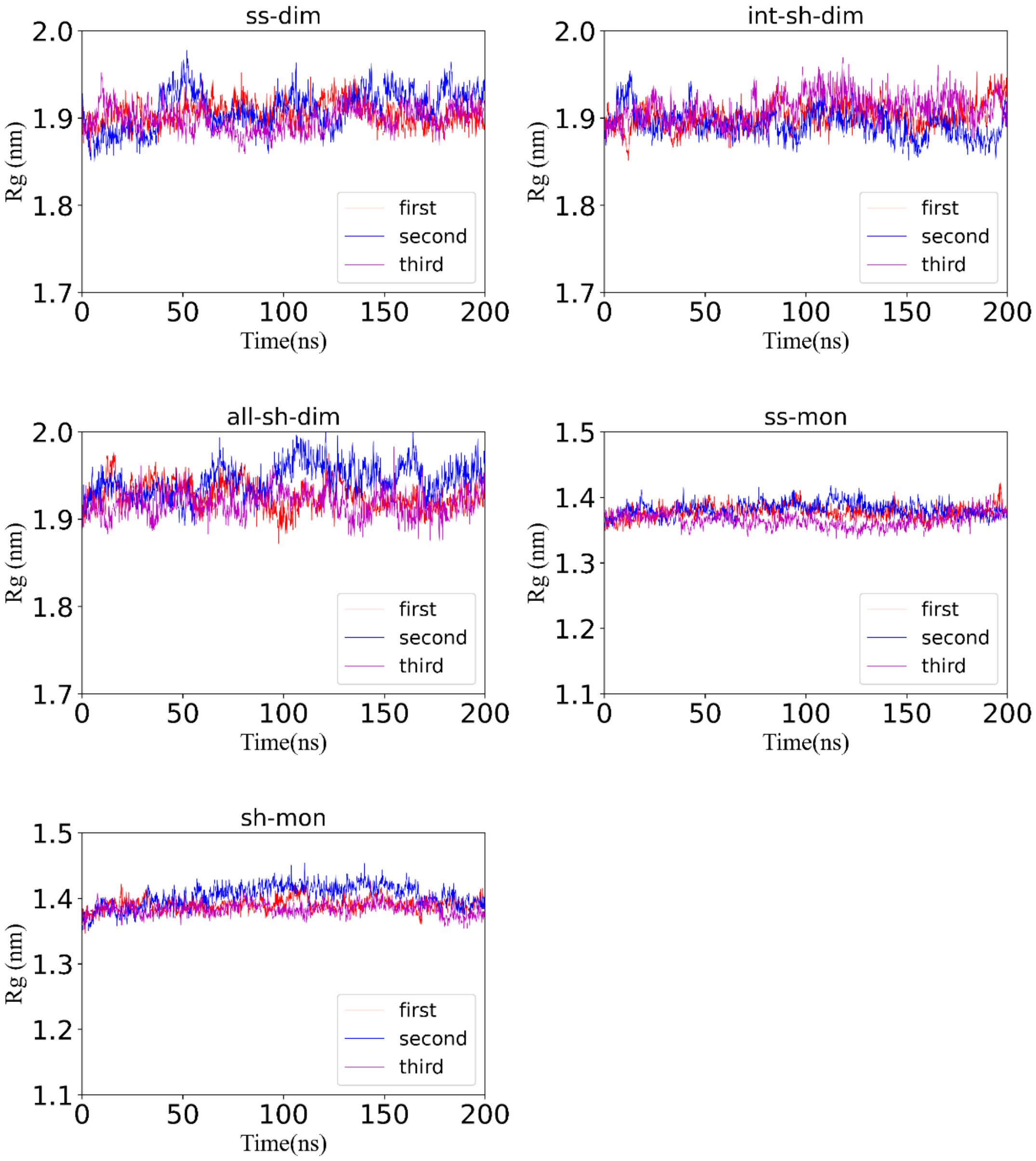


Figure S2. The Rg curves for the three replicated MD simulations on five different disulfide bonding states.

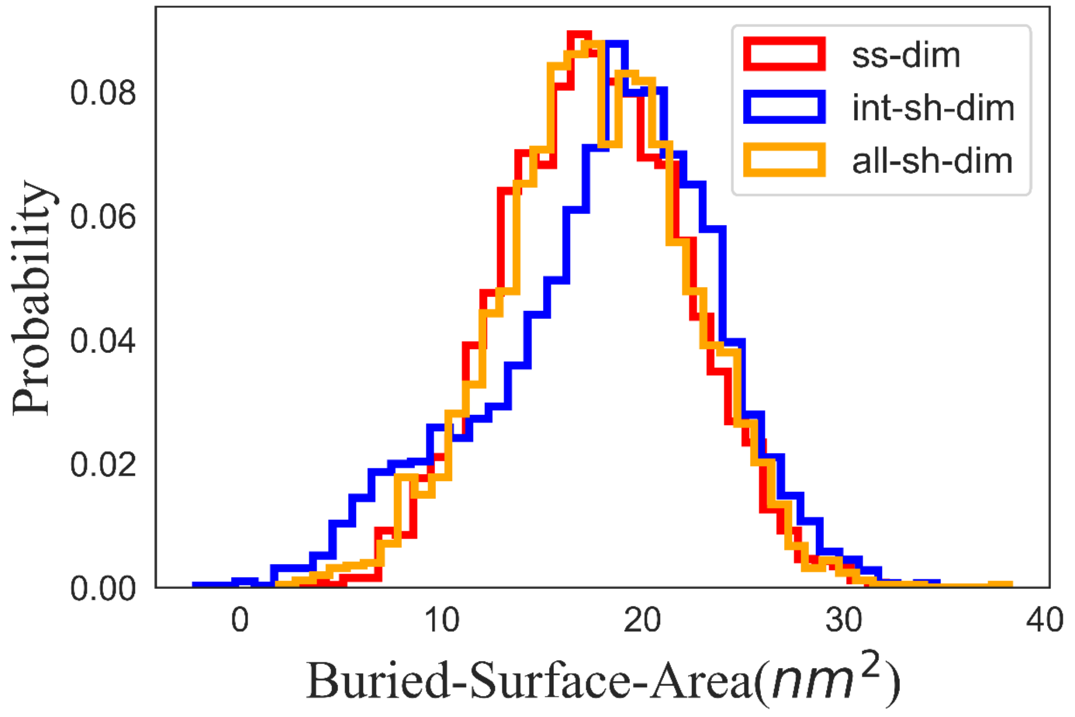


Figure S3. The distributions of the buried surface area for the three dimeric states.

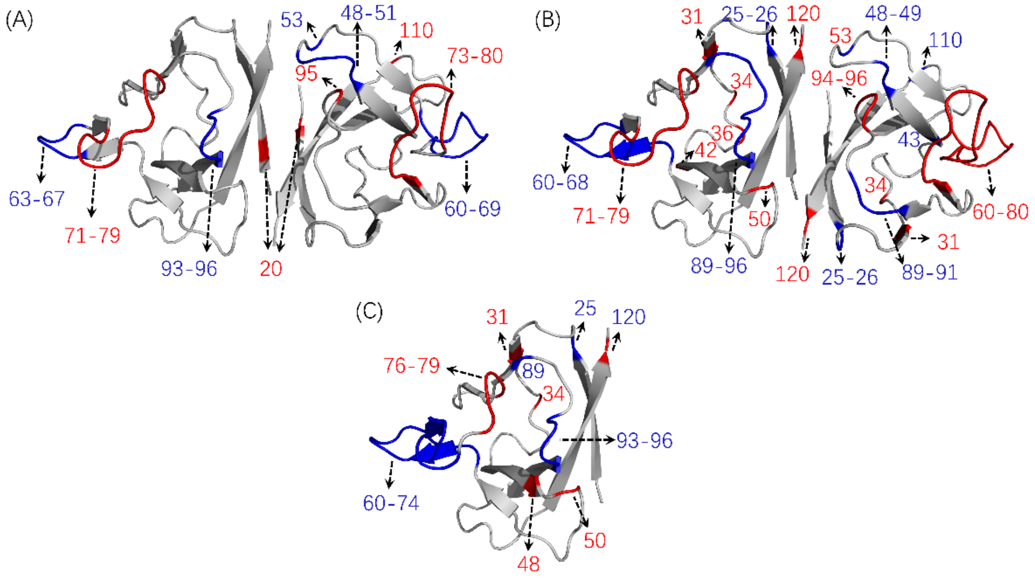


Figure S4. The locations of the residues that most affected by different disulfide bonds reduction schemes. (A) intermolecular disulfide bond reduction in dimer (B) all disulfide bonds reduction in dimer (C) intramolecular (all) disulfide bonds reduction in monomer.

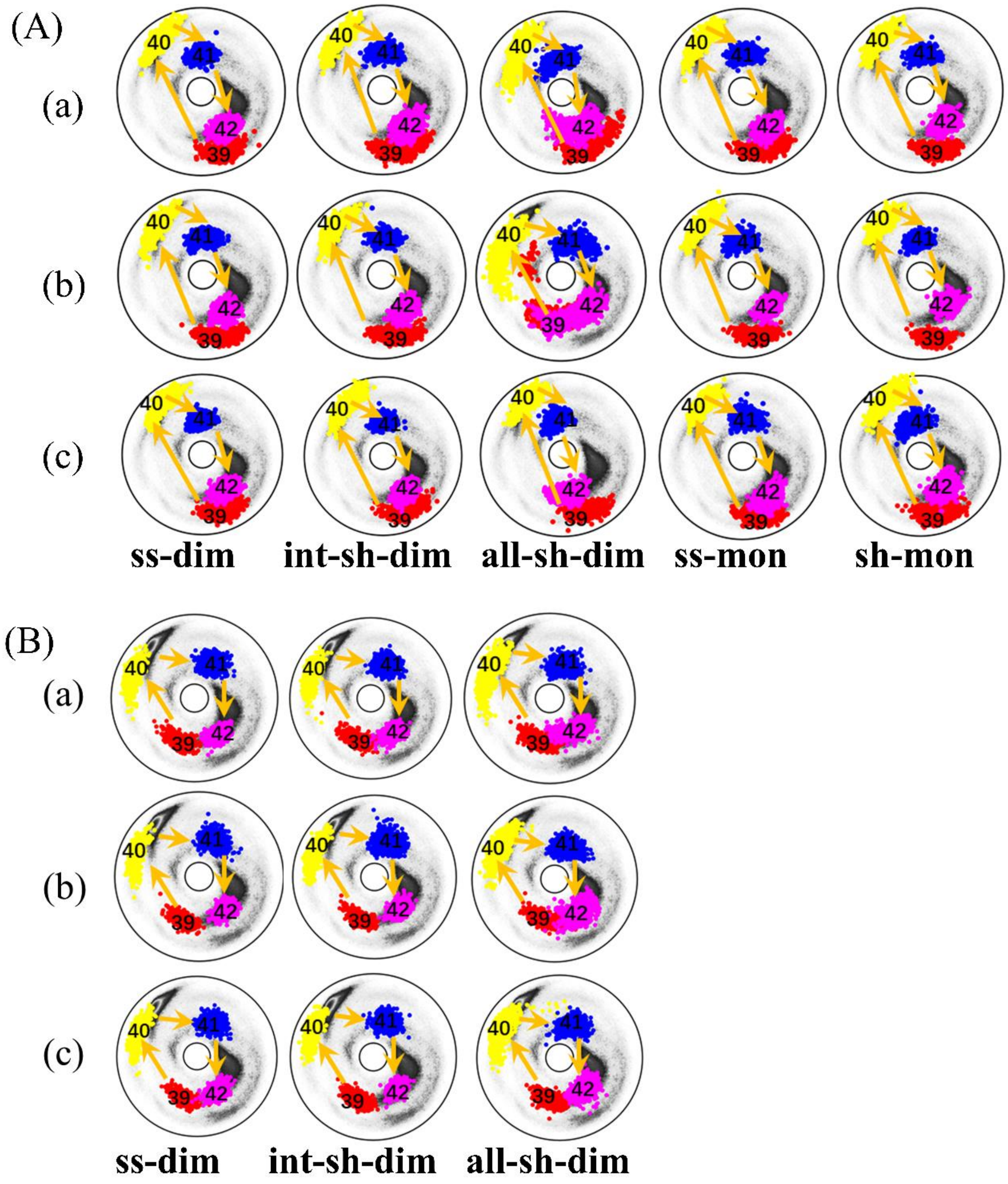


Figure S5. The distributions of the angle pairs ($\kappa_i$ ,$\tau_i$) with $i \in [39, 42]$ for the segment $^{39}$IHFY$^{42}$. Panel (A) and (B) are for chain A and B, respectively. In each panel, rows (a), (b), and (c) are the results from the first, second and third replicated simulation, respectively.

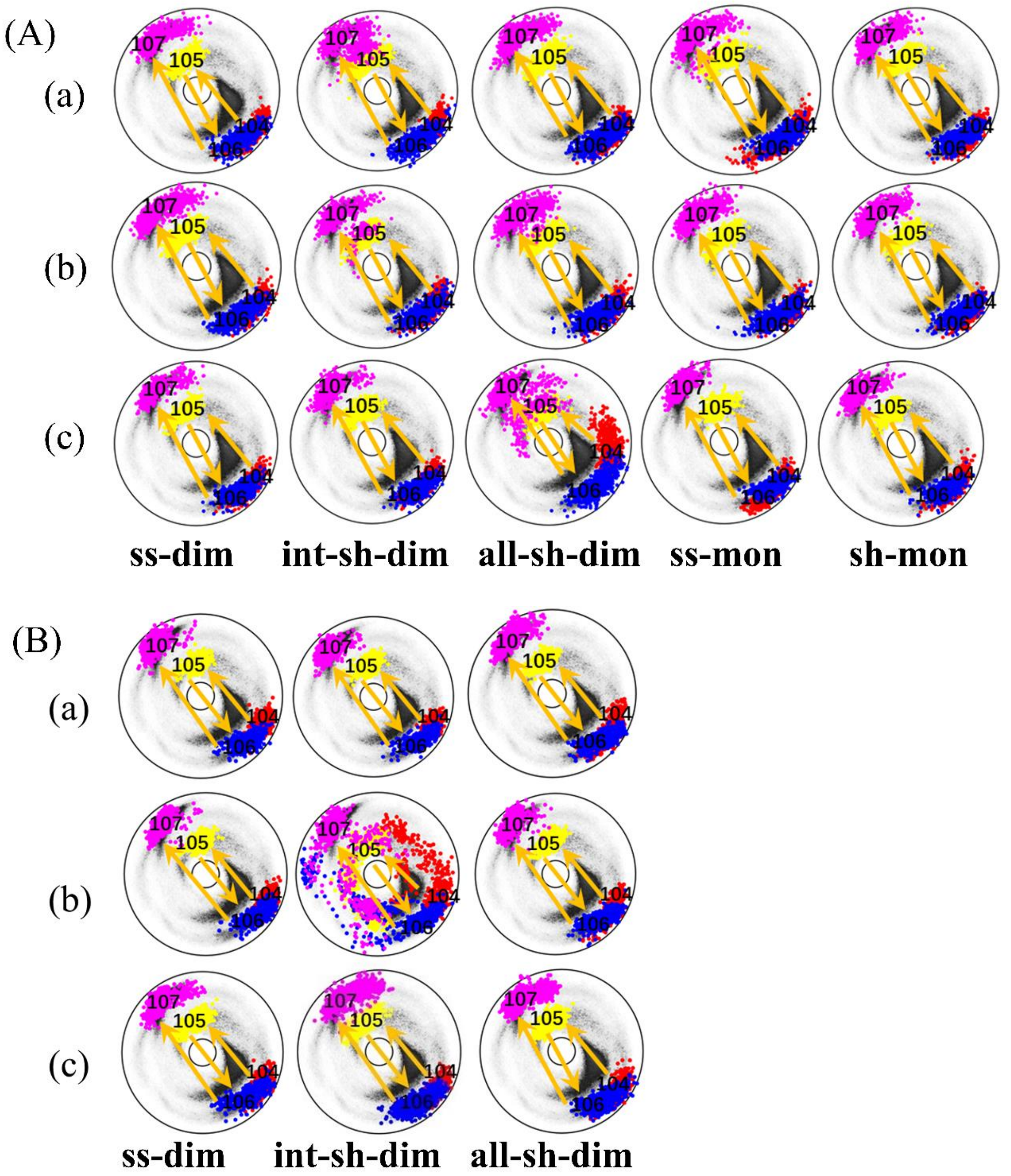


Figure S6. The distributions of the angle pairs ($\kappa_i$ ,$\tau_i$) with $i \in [104, 107]$ for the segment $^{104}$FYED$^{107}$. Panel (A) and (B) are for chain A and B, respectively. In each panel, rows (a), (b), and (c) are the results from the first, second and third replicated simulation, respectively.

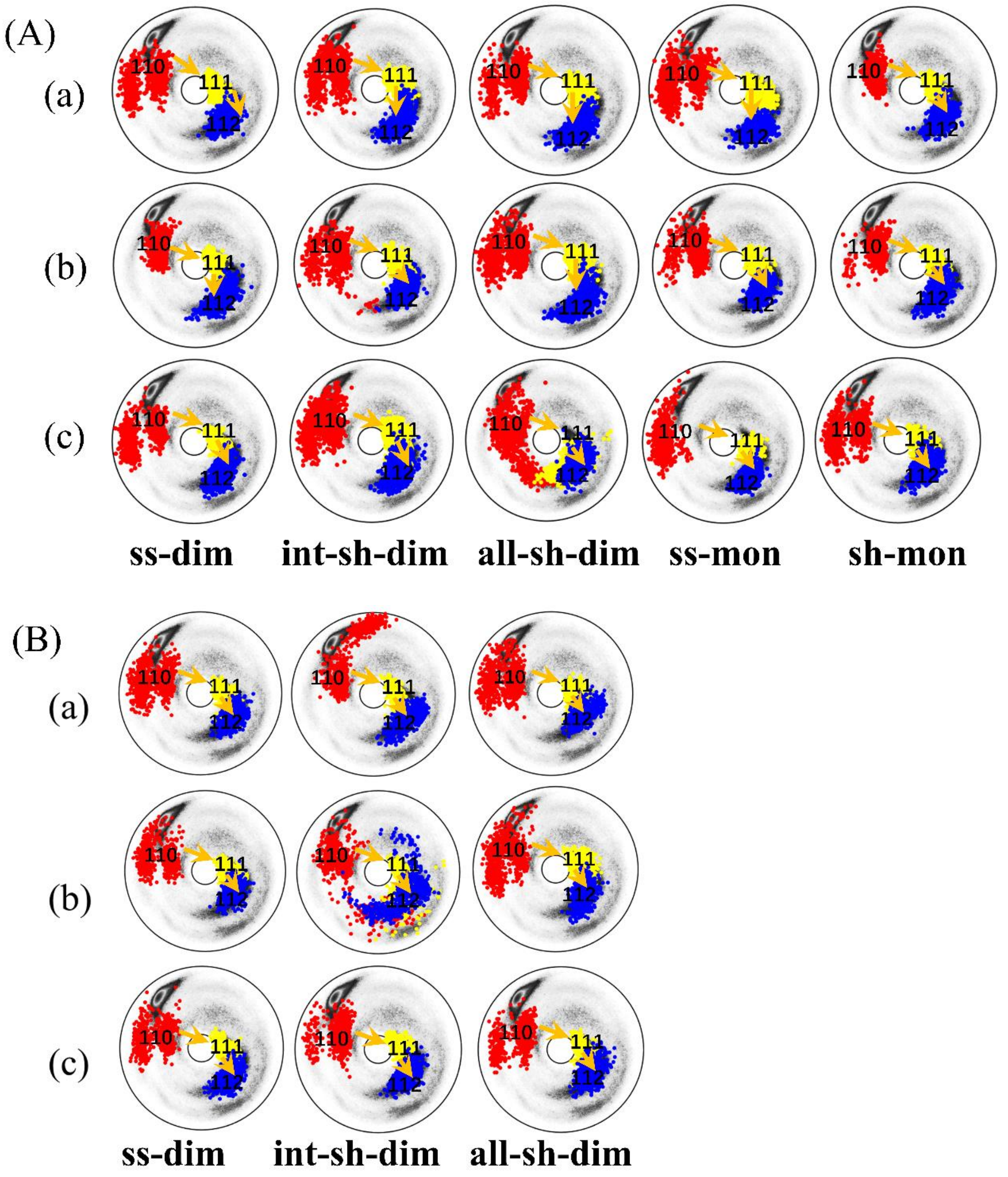


Figure S7. The distributions of the angle pairs ($\kappa_i$ ,$\tau_i$) with $i \in [110, 112]$ for the segment $^{110}$EYH$^{112}$. Panel (A) and (B) are for chain A and B, respectively. In each panel, rows (a), (b), and (c) are the results from the first, second and third replicated simulation, respectively.

(A)

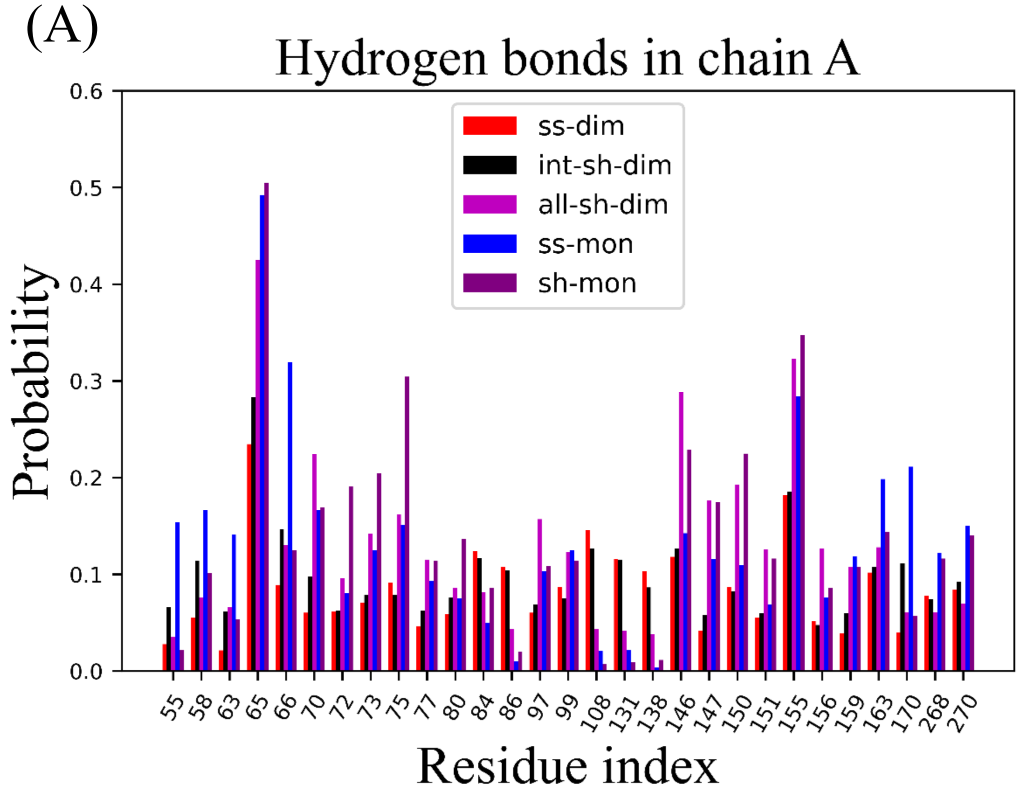


(B)

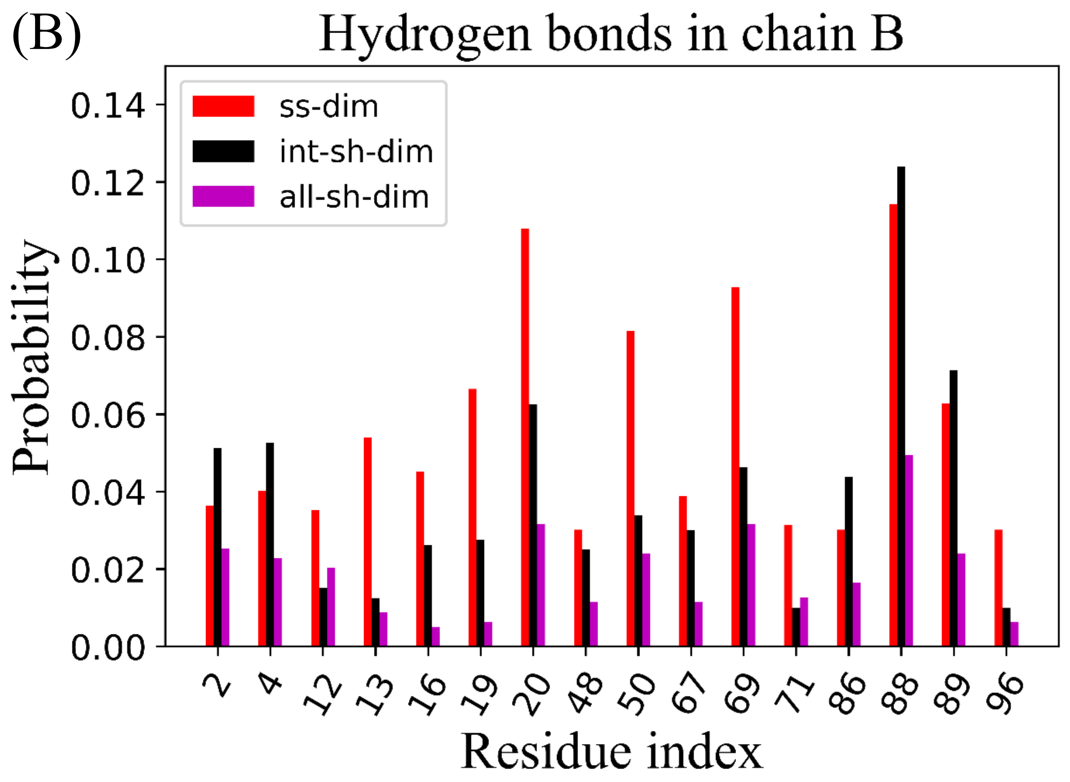


Figure S8: The distributions of the binding sites on HLA-A forming in hydrogen bonds with ORF8. (A) For chain A of HLA-A (B) For chain B of HLA-A.

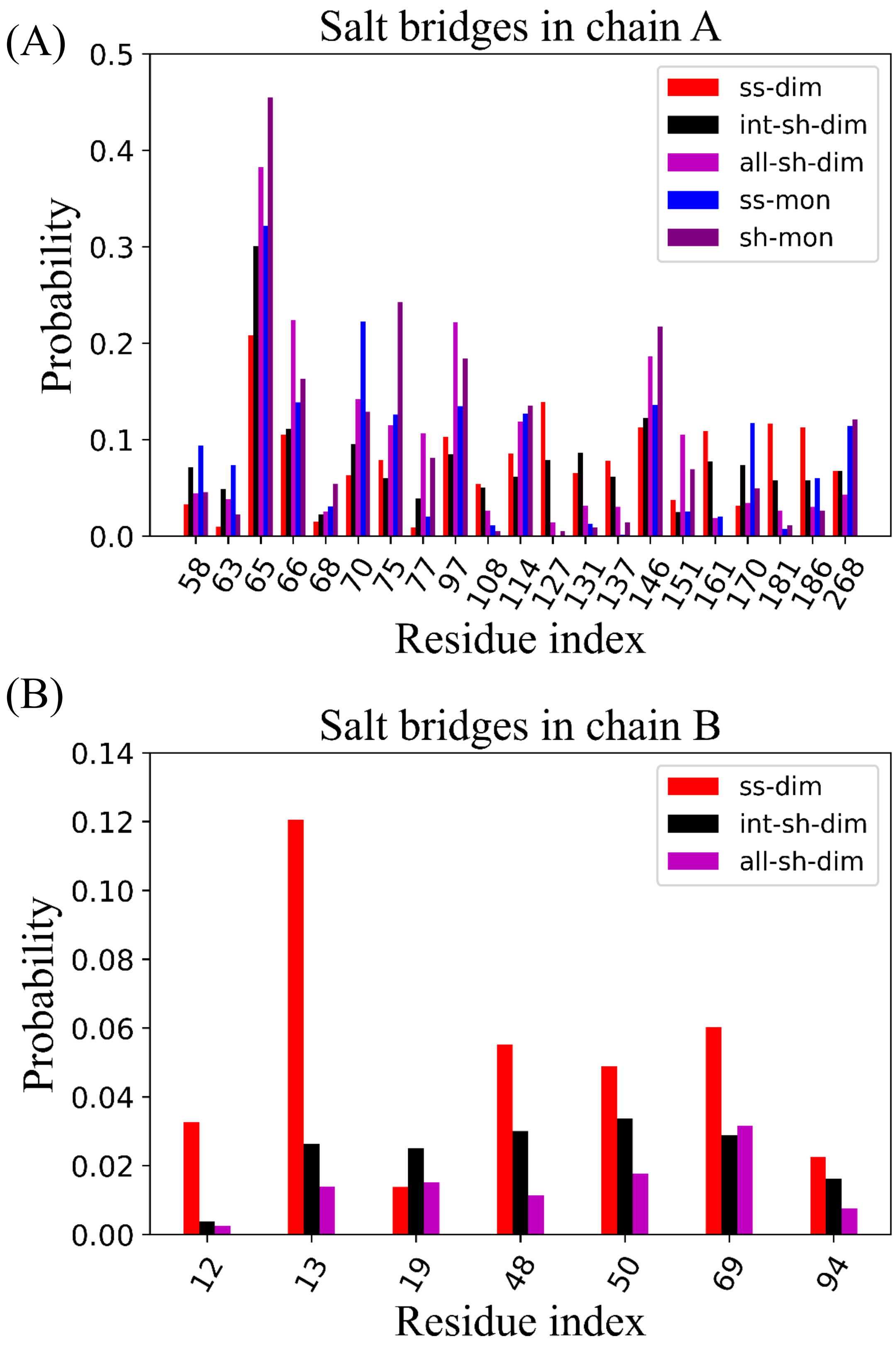


Figure S9. The distributions of the binding sites on HLA-A forming in salt bridges with ORF8. (A) For chain A of HLA-A (B) For chain B of HLA-A.

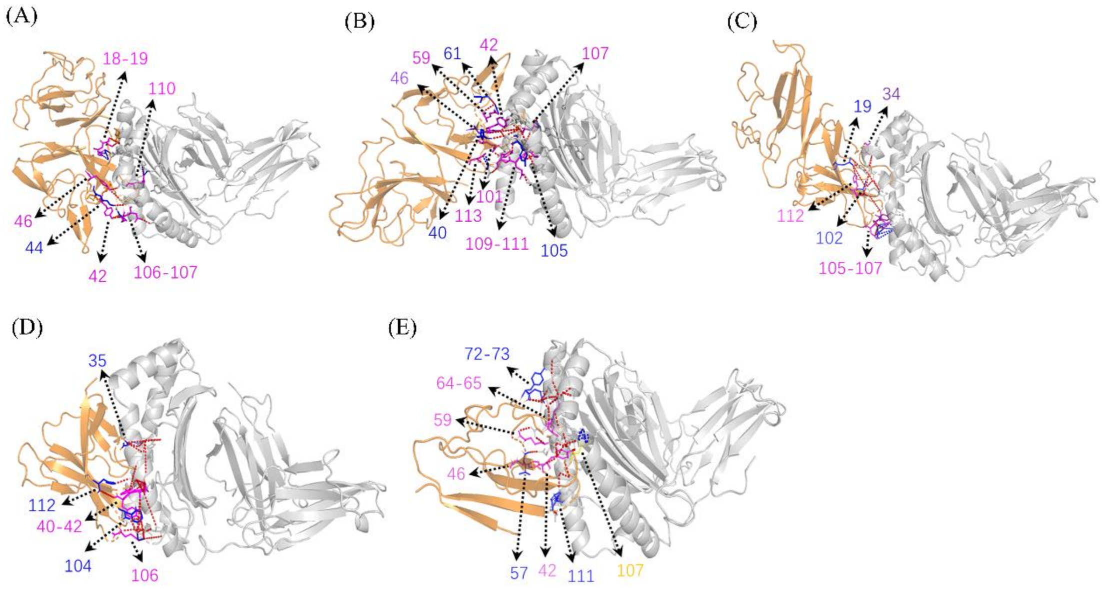


Figure S10. Five representative structures for the complex formed by ORF8 binding with HLA-A. The structure in gray is HLA-A, and the structure in orange is ORF8. The red dashed line represents the hydrogen bonds, and the blue dashed line represents the salt bridges. The residue on ORF8 involved in the formation of hydrogen bonds and salt bridges have been labeled. (A)ss-dim.(B)int-sh-dim.(C)all-sh-dim.(D)ss-mon.(E)sh-mon.

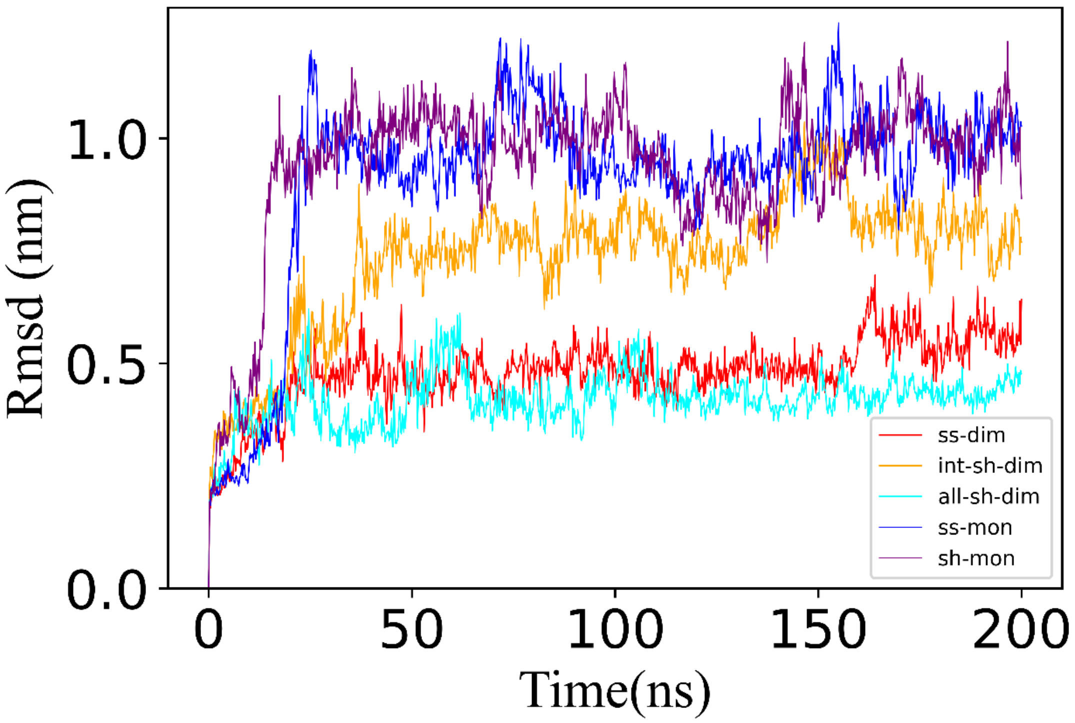


Figure S11. The curves of the RMSD evolution for the MD simulations on the five representative complexes.

.
